# On the relationship between indenation hardness and modulus, and the damage resistance of biological materials

**DOI:** 10.1101/107284

**Authors:** David Labonte, Anne-Kristin Lenz, Michelle L. Oyen

## Abstract

The remarkable mechanical performance of biological materials is based on intricate structure-function relationships. Nanoindentation has become the primary tool for characterising biological materials, as it allows to relate structural changes to variations in mechanical properties on small scales. However, the respective theoretical background and associated interpretation of the parameters measured via indentation derives largely from research on ‘traditional’ engineering materials such as metals or ceramics. Here, we discuss the functional relevance of indentation hardness in biological materials by presenting a meta-analysis of its relationship with indentation modulus. Across seven orders of magnitude, indentation hardness was directly proportional to indentation modulus, illustrating that hardness is not an independent material property. Using a lumped parameter model to deconvolute indentation hardness into components arising from reversible and irreversible deformation, we establish criteria which allow to interpret differences in indentation hardness across or within biological materials. The ratio between hardness and modulus arises as a key parameter, which is a proxy for the ratio between irreversible and reversible deformation during indentation, and the material’s yield strength. Indentation hardness generally increases upon material dehydration, however to a larger extend than expected from accompanying changes in indentation modulus, indicating that water acts as a ‘plasticiser’. A detailed discussion of the role of indentation hardness, modulus and toughness in damage control during sharp or blunt indentation yields comprehensive guidelines for a performance-based ranking of biological materials, and suggests that quasi-plastic deformation is a frequent yet poorly understood damage mode, highlighting an important area of future research.

## Introduction

In search for inspiration for the design of novel materials with enhanced properties, biological materials receive an increasing amount of attention (e. g. 1–5). Two features of these materials stand in stark contrast to ‘classic’ engineering materials: first, they are hierarchical in design, that is they are constructed from multiple building blocks arranged on characteristic length scales ranging from a couple of nanometres to a few millimetres. Second, the structural arrangement and properties of these components are tailored to specific functional demands.

A thorough understanding of how the arrangement and properties of individual components determine the performance of the ‘bulk’ materials is a prerequisite for any attempt at replicating their functionality. Hence, mechanical and structural characterisation methods traditionally developed for and applied to engineering materials are increasingly used to study biological materials. Among these, instrumented indentation has been particularly popular, due to the relative ease of use and its ability to accurately measure material properties on various length scales (6–8). In biological materials, instrumented indentation is frequently used as a tool to relate structural changes, for example degree of mineralisation, to variations in material properties, such as fracture toughness, elastic modulus, and hardness. These properties, in turn, determine performance and hence the adaptive value of the structural changes in question, and therefore can shed light on critical structurefunction relationships.

Fracture toughness, *T*, quantifies a material’s resistance to crack initiation and propagation, and is superseding ‘strength’ as the property most associated with high-performance materials (9, 10). The hierarchical design and the ensuing complexity of biological materials necessitates a careful execution and application of the existing indentation methodology (11). However, instrumented indentation remains an unrivalled method for the evaluation of toughness, in particular in biological materials which are frequently too small or irregular to allow the fabrication of ‘standardised’ specimen required for more conventional fracture tests (8).

The elastic modulus quantifies the resistance to elastic (reversible) deformation. In contrast to toughness, a large modulus is not necessarily desired (for example, adhesive structures need to be somewhat compliant), so that a direct link between its magnitude and performance is often not straightforward. Most studies assume isotropic behaviour of biological materials, i. e. a mechanical response that is independent of the testing direction. However, it is frequently acknowledged that this assumption is more practical than accurate, and most of the literature hence reports an indentation modulus, *E*_I_, which is equal to the plain strain modulus, *E′*, if the modulus of the indenter is much larger than that of the material, and if the material is isotropic.

Hardness is probably the most commonly measured, but least understood of the material properties that can be assessed by instrumented indentation. Despite an increasing body of literature which has highlighted that hardness is a complex property that needs to be interpreted with care (see e. g. 6, 12–17), it is still often taken as a proxy for the resistance to ‘plastic’ (i. e. irreversible) deformation. This interpretation has been particularly popular in the more biological literature, perhaps because it provides a straightforward link to functional and hence ecological relevance. Hardness has long been a property of considerable importance in engineering materials: it is an index of strength (18) and – in combination with toughness – of brittleness (19). Hardness is frequently associated with a material’s resistance against being penetrated (i. e. elastically deformed), spread (i. e. irreversibly deformed) or scratched (i. e. fractured) (20). Clearly, the physical mechanisms involved in these processes differ. What is the functional meaning of indentation hardness?

In this article, this question is addressed by presenting a comprehensive meta-analysis of published data on indentation modulus, *E*_I_, and indentation hardness, *H*_I_, of diverse biological materials. We investigate the relationship between the two properties, which is then interpreted with a series elastic-plastic deformation model. This analysis provides guidelines for a careful interpretation and comparison of indentation hardness across and within biological materials, and highlights potential pitfalls. Next, we focus more specifically on the functional relevance of *T*, *E*_I_ and *H*_I_ for the avoidance and containment of mechanical damage introduced by ‘blunt’ or ‘sharp’ contacts,^1^ and discuss the competition between quasi-plasticity and fracture in biological materials. Quasi-plasticity emerges as the dominant damage mode for most biological materials, which is controlled not by indentation hardness, but by its ratio with the indentation modulus.

## Analysis & Discussion

### The correlation between indentation hardness and modulus

Indentation hardness and modulus of various biological materials were assembled from the literature (22– 89, all data are available in the supplemental material). Across seven orders of magnitude, indentation hardness was directly proportional to indentation modulus (figure 1), with a proportionality constant of approximately 0.05, 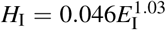 (reduced major axis regression on log-transformed data, 95% confidence intervals (CI) [0.043; 0.050] and [1.01; 1.06]). The strength of the correlation, R^2^ = 0.96, is somewhat deceptive, as the regression was performed on log-transformed data. Some of the variation in the ratio *H*_I_/*E*_I_ across well-represented materials is captured in Table 1. The data used in the regression include properties measured with different indenter geometries, and in hydrated or dehydrated conditions. These factors are discussed further below.

**Figure 1.**
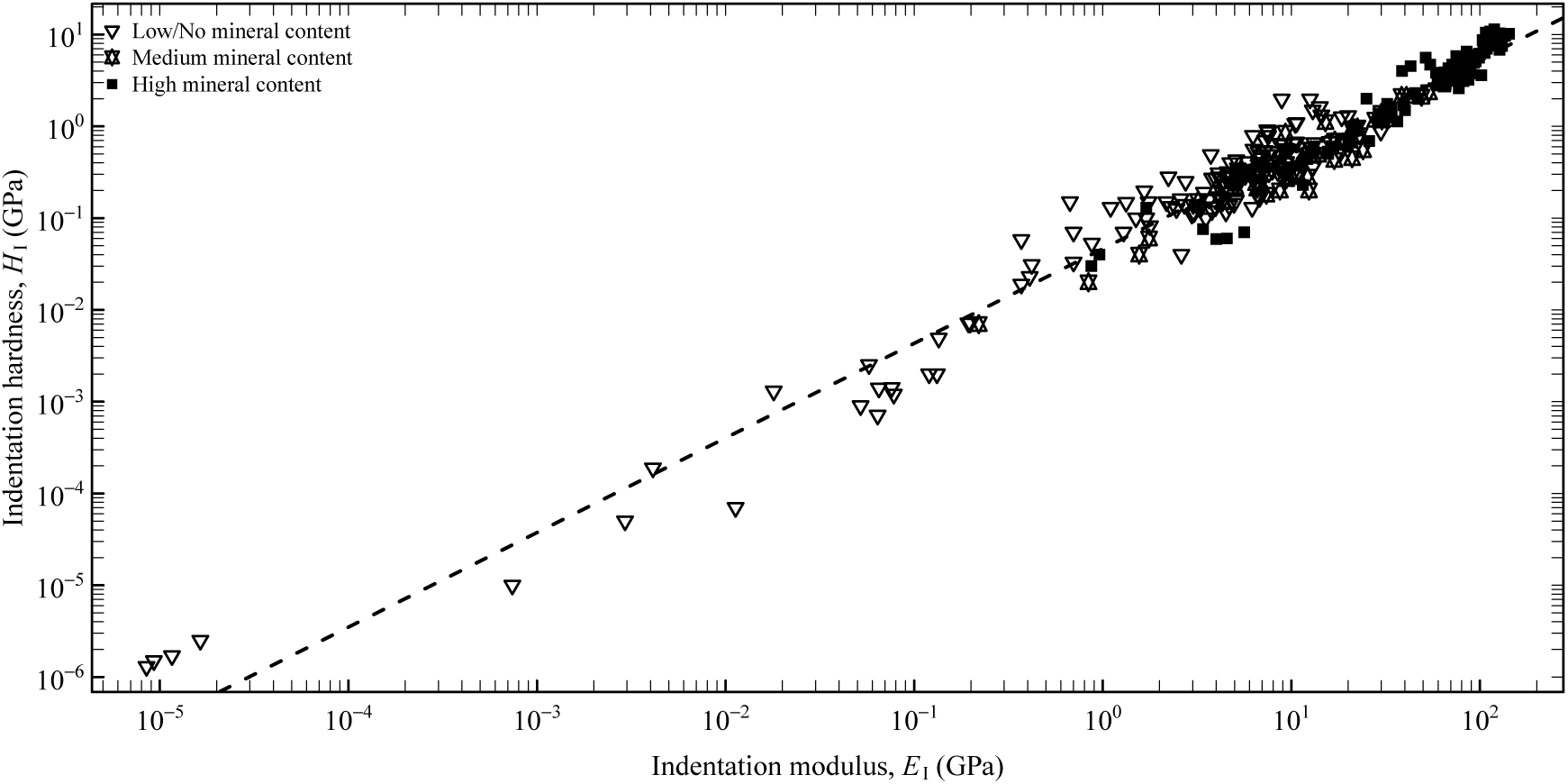
Across seven orders of magnitude, indentation hardness, *H*_I_, was directly proportional to indentation modulus, *E*_I_. The dashed line is the result of a reduced major axis regression on log-transformed values, 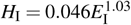 (95% confidence intervals (CI) [0.043; 0.050] and [1.01; 1.06], R^2^ = 0.96). An ordinary least-squares regression yielded a similar result, 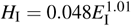 (95% CI [0.045; 0.052] and [0.99; 1.03]. Thus, *H*_I_ ≈ 0.05*E*_I_. Classification by mineral content was performed as described in Amini & Miserez, 2013 (21).

**Table 1:** While indentation hardness, *H*_I_, and indentation modulus, *E*_I_, were linearly related across seven orders of magnitude, their ratio, the elasticity index *I*_E_, differed systematically across different materials, indicating different amounts of relative irreversible deformation, *h*_r_/*h*_max_ (calculated using the median values for *I*_E_ and eq. 3). Data are ordered by their elasticity index.

| Material | N | Indentation modulus, $E_I$ (GPa) | | Indentation hardness, $H_I$ (GPa) | | $I_E$ | $h_r/h_{\max}$ (%) |
| --- | --- | --- | --- | --- | --- | --- | --- |
|  |  | Median | Range | Median | Range | Median |  |
| Crustacean Cuticle | 15 | 19.0 | 3.2-53.7 | 0.45 | 0.15-2.35 | 0.033 | 0.82 |
| Dentin | 10 | 26.5 | 8.7-49.0 | 1.00 | 0.20-2.10 | 0.040 | 0.78 |
| β-Keratin | 11 | 4.3 | 3.2-6.0 | 0.19 | 0.13-0.41 | 0.044 | 0.75 |
| Mineralised with Calcite | 16 | 65.7 | 25.9-73.0 | 3.35 | 0.69-4.70 | 0.052 | 0.71 |
| Insect Cuticle | 67 | 6.7 | 0.4-29.8 | 0.33 | 0.03-1.98 | 0.054 | 0.70 |
| Enamel/Enameloid | 16 | 96.0 | 62.7-131.2 | 5.90 | 2.60-7.60 | 0.057 | 0.68 |
| Wood | 14 | 5.9 | 1.1-18.1 | 0.33 | 0.13-0.41 | 0.063 | 0.65 |
| Mineralised with Aragonite | 30 | 90.6 | 36.5-142.5 | 5.40 | 1.13-11.40 | 0.068 | 0.62 |

A correlation between hardness and bulk, Young’s and in particular shear modulus has also been reported for metals, bulk metallic glasses, ceramics and polymers (see e. g. 90–99), which can be qualitatively understood by considering two limiting cases: In rigid-perfectly-plastic materials, all deformation is accommodated irreversibly, and hence no *elastic* energy is stored in the deformed material. In this limit, indentation hardness is indeed an accurate proxy for the resistance to irreversible deformation. If the yield stress is infinite, in turn, all deformation is accommodated reversibly, so that the mean contact pressure – and thus the indentation hardness – is entirely determined by the elastic properties of the material and the indenter geometry. The reality lies in between these two cases, so that indentation hardness is hardly ever independent of indentation modulus (but not *vice versa*. Notably, this problem cannot be circumvented by measuring the lateral extent of the residual impression as is done in ‘traditional’ hardness tests, because the elastic recovery occurs often dominantly in-depth, see e. g. 100, 101). The indentation hardness is determined by properties pertaining to the resistance against both irreversible and reversible deformation, and exploring its correlation with indentation modulus further therefore requires a mechanical model that accounts for both.

Using the classic framework of lumped parameter models, irreversible and reversible deformation may be assumed to be linked in series, as utilised by Sakai (14). In this simple model, the elastic component is characterised by an elastic modulus (represented by a spring), while the irreversible component is characterised by a ‘true’ resistance to irreversible deformation, *H*, which is equivalent to the mean contact pressure during the indentation of a rigid-perfectly-plastic material (represented by a slider). The respective constitutive relationships for perfectly plastic and purely elastic conical indentation can be combined to obtain (a detailed derivation can be found in the supplemental material, 14, 17, 102)):

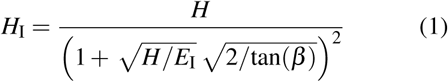

or equivalently

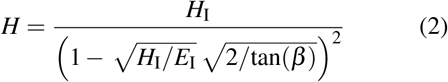

where *β* is the equivalent cone angle of the indenter. A number of conclusions are immediately apparent from these equations:

1. *As H*_I_*/E*_I_ → 0, the indentation hardness converges to the ‘true’ hardness, i. e. *H/H*_I_ → 1 (figure 2 A). Hence, indentation tests reasonably approximate resistance to irreversible deformation in materials with small values of *H*_I_*/E*_I_, for which they were originally developed and remain best-understood. For example, for ductile metals *H*_I_*/E*_I_ ≈ 10^−3^, and hence the indentation hardness measured with Vickers or Berkovich tips is within 20% of the ‘true’ hardness.
2. *As H*_I_*/E*_I_ → 1/2tan(*β*), *the true hardness diverges,* because the indentation hardness cannot exceed the elastic contact pressure, *H*_I_ ≤ 1/2*E*_I_tan(*β*) (see figure 2 A). We found several reports in contradiction with this limit (e. g. in 42, 53, 83), most likely explained by viscoelastic behaviour of the indented materials.
3. *The indentation hardness depends on the indenter geometry.* The influence of the effective angle is negligible when data obtained with Vickers, Berkovich or Knoop indentors are compared, but can be significant when results from indentations with cube corner or spherical indenters are included (a more detailed analysis of this dependence can be found in reference 14). We used eqs. 1 and 2 to rescale the data shown in figure 1 to Berkovich indentation hardness values, and found that the estimated relationship remained virtually unchanged, illustrating the robustness of the collated data (see supplemental material).
4. *The ‘true’ resistance to plastic deformation, H, depends on the* ratio *between indentation hardness and indentation modulus.* Thus, a large indentation hardness does not imply a large resistance to irreversible deformation *per se*. As an illustrative example, the mandibles of West Indian dry-wood termites (*Cryptotermes brevis*) have a Berkovich indention hardness of 1.07 GPa, and an indentation modulus of 10.5 GPa (34), implying a ‘true’ hardness of 17.8 GPa. Hydrated human enamel, in turn, has a Berkovich indentation hardness which is thrice as large (3.21 GPa), but its ‘true’ hardness is almost two times smaller (*H* = 10.8 Gpa), because the indentation hardness is dominated by the large indentation modulus of 87.02 GPa (35).

**Figure 2.**
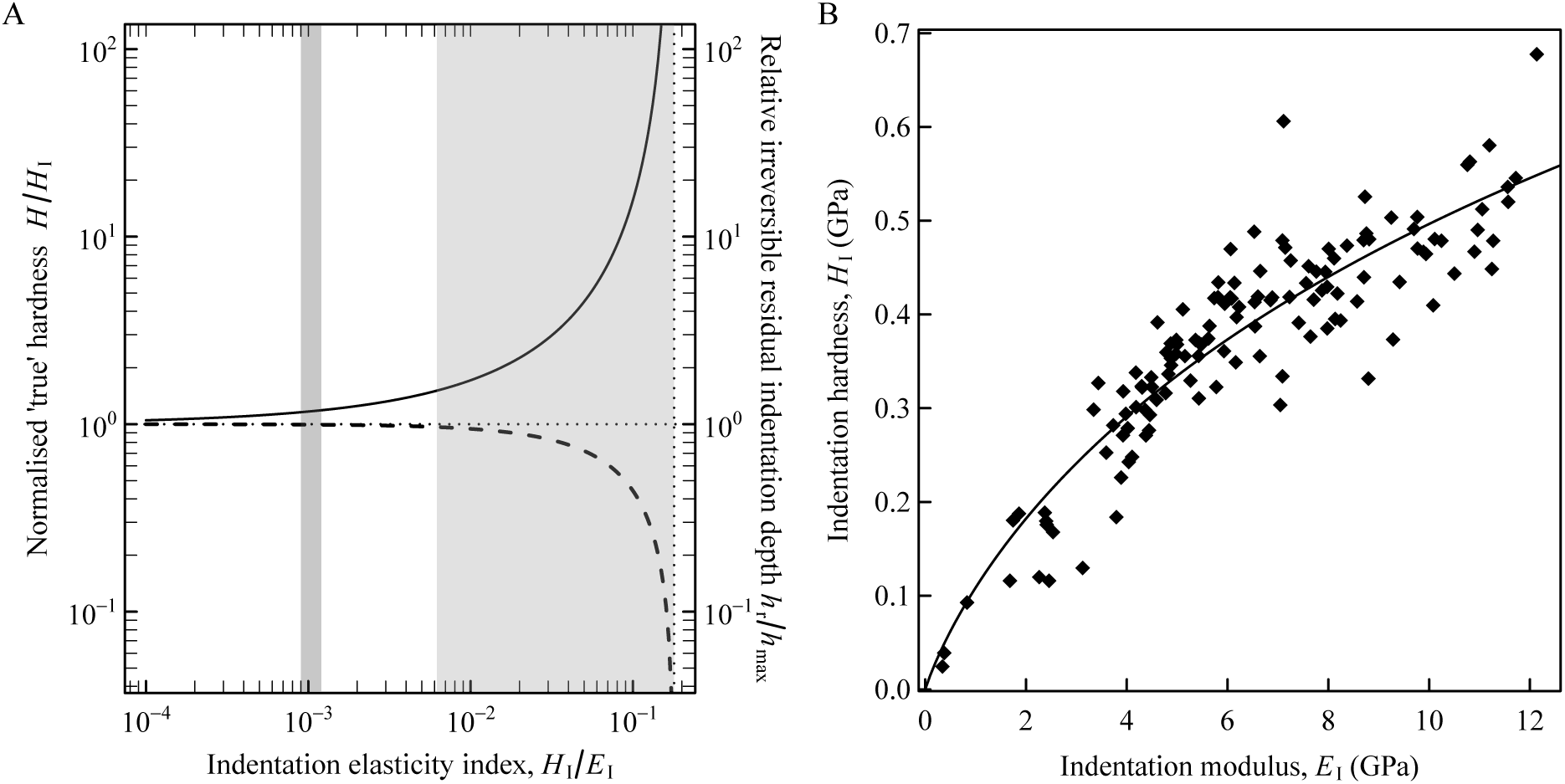
(A) The ‘true’ indentation hardness, *H*, can be estimated from the indentation elasticity index, *I*_E_ = *H*_I_*/E*_I_, by assuming that reversible and irreversible deformation are linked in series (see text). The ratio *H/H*_I_ (solid line, eq. 2 with *β* = 19.7^*◦*^ for a Berkovich indenter) approaches 1 in the limit of small *I*_E_, while it diverges as *I*_E_ approaches its maximum possible value set by the elastic contact pressure, *I*_E_ ≤ tan(*β*)/2 ≈ 0.18 for a Berkovich indenter. As a consequence, the deformation transitions from dominantly irreversible for materials with small values of *I*_E_ to effectively completely reversible for large values of *I*_E_, as illustrated by the dashed line (eq. 3). The dark grey rectangle illustrates the range of *I*_E_ typical for ductile metals, in which *H/H*_I_ is close to one; the light grey rectangle indicates the range observed for biological materials. (B) As indentation hardness is a hybrid measure related to both irreversible and reversible deformation, changes in indentation modulus alone will change the measured indentation hardness. The adhesive setae of Chilean rose tarantulas were indented at different angles to their axis in order to study their anisotropy (103). The observed almost 30-fold change in indentation hardness is likely solely based on a change of the indentation modulus, as indicated by a non-linear least-squares fit of eq. 1 (solid line), which yielded a constant ‘true’ hardness of 2.21 GPa (95% confidence interval (2.12; 2.33)).

The correlation between indentation hardness and indentation modulus requires particular attention in materials with pronounced anisotropy or gradual changes in material properties. In order to illustrate this point, we briefly discuss the relationship between indentation hardness and modulus in the adhesive hairs (setae) of Chilean rose tarantulas (*Grammostola rosea*), indented at different angles relative to their longitudinal axis (103). While *H*_I_ varied by a factor of almost 30 (figure 2 B), a non-linear least-squares fit of eq. 1 suggested that much of this variation is explained by a change of the indentation modulus – most likely caused by the unidirectional orientation of reinforcing fibres in the adhesive setae themselves – and that the ‘true’ hardness is actually constant at 2.21 GPa (95% confidence interval (2.12; 2.33)).

The previous discussion underlines that any interpretation of changes in *H*_I_ within or across structural biological materials requires care, and should at the very least include the ratio *H*_I_*/E*_I_. This ratio can also be used to calculate a ‘true’ hardness, a property independent of the modulus and indenter geometry, which represents the energy required to create a unit volume of irreversible impression (14), and thus provides additional comparative information about the mechanical performance of biological materials. The mechanistic origin of the increase of *H* with *E*_I_ implied by the available data requires further clarification.

The verdict that indentation hardness is a hybrid measure characterising reversible and irreversible deformation is not new (see e. g. 6, 12–16), but has received relatively little attention outside the more technically oriented indentation literature. A related interpretation is that *H*_I_ is a measure for the work required to create a unit volume of residual impression (14), and that the ratio *H*_I_*/E*_I_ is thus proportional to the ratio between the irreversible and reversible work done during indentation (13, 14). As a consequence, *H*_I_*/E*_I_ may be interpreted as an ‘elasticity index’ *I*_E_ = *H*_I_*/E*_I_: Large values indicate a dominance of reversible deformation, and small values imply dominantly irreversible deformation (14). The elasticity of the contact can also be assessed from the slopes of the loading and unloading curves (104), or from the ratio between the residual indentation depth, *h*_r_, and the indentation depth at maximum load, *h*_max_ (105). In the framework of the Oliver-Pharr analysis, this ratio is given by:

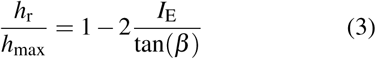

The direct proportionality between *H*_I_ and *E*_I_ across seven orders of magnitude implies that the elasticity index is constant, *I*_E_ ≈ 0.05, and hence that around 3/4 of the total deformation of biological materials during indentation with a Berkovich indenter is irreversible (*h*_r_/*h*_max_ *≈* 72%, see figure 2 A).

### The effect of hydration on indentation hardness and modulus

There is ample evidence that the mechanical properties of biological materials are sensitive to hydration, which tends to decrease hardness and modulus, but increase toughness (e. g. 3, 22, 35, 60, 71, 106– 109). It is convenient to define the sensitivity to hydration, *S*_H_, as the relative change in indentation hardness/modulus upon dehydration, *S*_H,*H*_I__ = (*H*_I, d_ − *H*_I, w_)*/H*_I, w_ and *S*_H,*E*_I__ = (*E*_I, d_ *− E*_I, w_)*/E*_I, w_, where the indices *w* and *d* indicate data from hydrated (‘wet’) and dehydrated (‘dry’) materials, respectively. Dehydration can affect the measured mechanical properties by a change in both structure and/or properties of the material, and hydrophobic and hydrophilic materials will take up different amounts of water, even if their ultrastructure is identical. The effects of these factors cannot be addressed with the presently available data.

A meta-analysis of data collated from the literature suggests that the sensitivity to hydration is largest for materials with small *H*_I, w_ and *E*_I, w_ (see figure 3). A regression of the combined data – from 0.007 GPa < *H*_I, w_ < 8.7 GPa and 0.22 GPa < *E*_I, w_ < 125.7 GPa – suggested that the sensitivity to hydration decreases with hydrated indentation hardness as 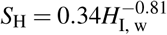 (R^2^ =71%, 95% CI (0.26; 0.42) and (-0.91; 0.71), respectively. An analogous result with respect to *E*_I, w_ is given in the supplemental material). Indentation properties of dehydrated biological materials hence have to be interpreted with great care: First, they will almost always significantly deviate from the ‘true’ value – even stiff and hard materials with *H*_I, w_ *>* 4.5 GPa (or *E*_I, w_ *>* 90 GPa) change their indentation properties by up to 10% upon dehydration. Second, while it may be possible to predict dehydrated from the hydrated properties, the inverse will be more error-prone.

**Figure 3.**
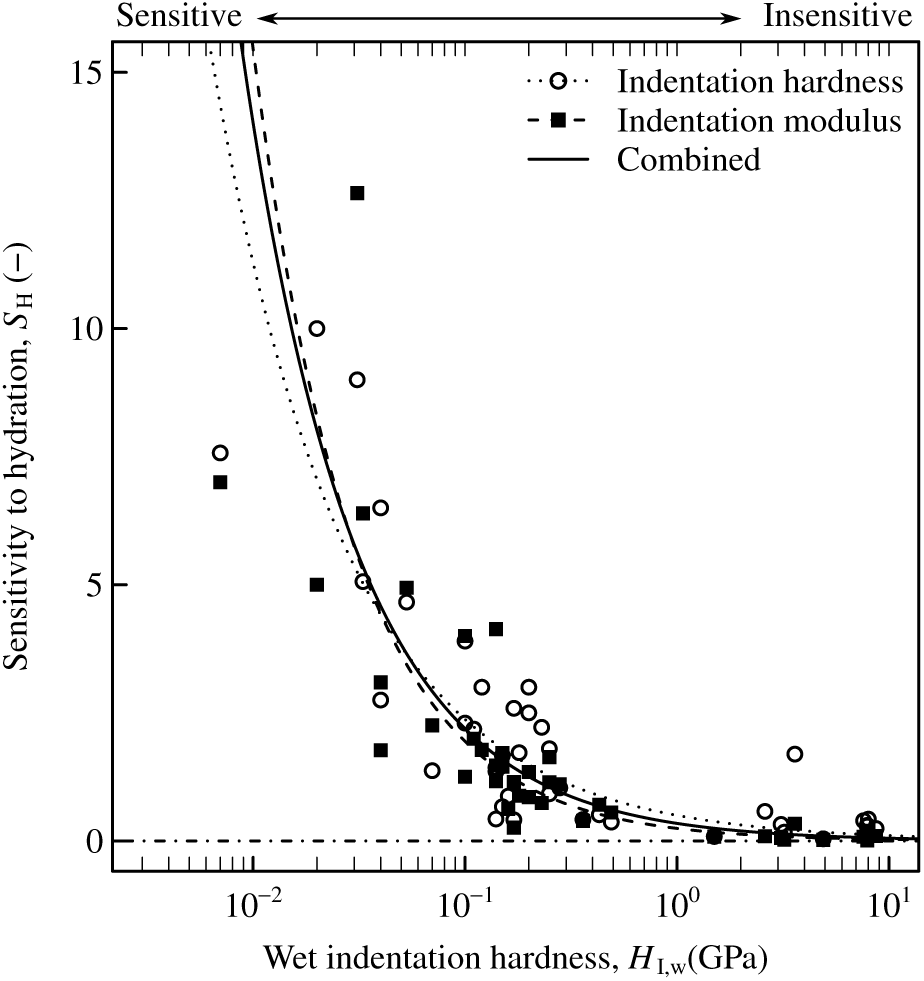
Biological materials are particularly sensitive to hydration if their hydrated indentation hardness or modulus is small (note the logarithmic x-axis). The sensitivity to hydration is defined in this context as (*H*_I, d_ – *H*_I, w_)*/H*_I, w_ and (*E*_I, d_ – *E*_I, w_)*/E*_I, w_, i. e. *S*_H_ = 0.2 indicates a 20% increase in hardness and modulus upon dehydration, whereas *S*_H_ = 0 indicates no change. The lines are the result of reduced major axis regression on log-transformed data, separately for indentation hardness (dotted line), indentation modulus (dashed line), and both combined (solid line). The detailed results of the regressions are given in the supplemental material.

As indention hardness is a hybrid measure of a material’s resistance to irreversible and reversible deformation, a change in indentation modulus will also affect indentation hardness. In order to investigate if the effect of hydration on indentation hardness is explained by the associated change in indentation modulus, we estimated the ‘true hardness’ of hydrated materials via eq. 2, and then predicted the indentation hardness of dry materials using eq. 1. This procedure resulted in a significant underestimation of *H*_I, d_ (by a factor of around 1.5, Wilcoxon signed-rank test, V = 1112, p = 0.012), suggesting that the ‘true’ resistance to irreversible deformation increased upon dehydration in most of the investigated materials. Hence, water generally acts as a ‘plasticizer’.

### Damage control in biological materials

In both biological and engineering contexts, indentation hardness is frequently used as a proxy for wear resistance, either directly, in ratio to indentation modulus, or in form of the property group 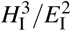 (e. g. 21, 34, 36, 110–113). The diverse ways in which hardness can be related to wear resistance partly reflects the fact that wear is a complex and system-dependent phenomenon (114). As a consequence, different wear mechanisms are controlled by different parameter groups (21, 115).

Resistance to mechanical wear quantifies the propensity of a material to be permanently altered (damaged) by mechanical interactions with external bodies/particles, for example via quasi-plastic deformation (yield) or fracture (cracking). These two phenomena are governed by different physical processes, so that the material properties that determine wear resistance depend on the dominating damage mode. As it is unlikely that damage can be avoided altogether (for example, surface craters and cracks are abundant in hominid teeth, see ref. 116), wear resistance becomes a question of damage containment. Both quasi-plasticity and fracture give rise to a damage zone with a characteristic length (schematically shown in figure 4), which can be used to rank materials by their damage resistance. For simplicity, we assume (i) that the wear contact is frictionless, and (ii) that the plane strain modulus of the abrasive material is much larger than that of the abraded material (for a discussion including friction, see ref. 115).

**Figure 4.**
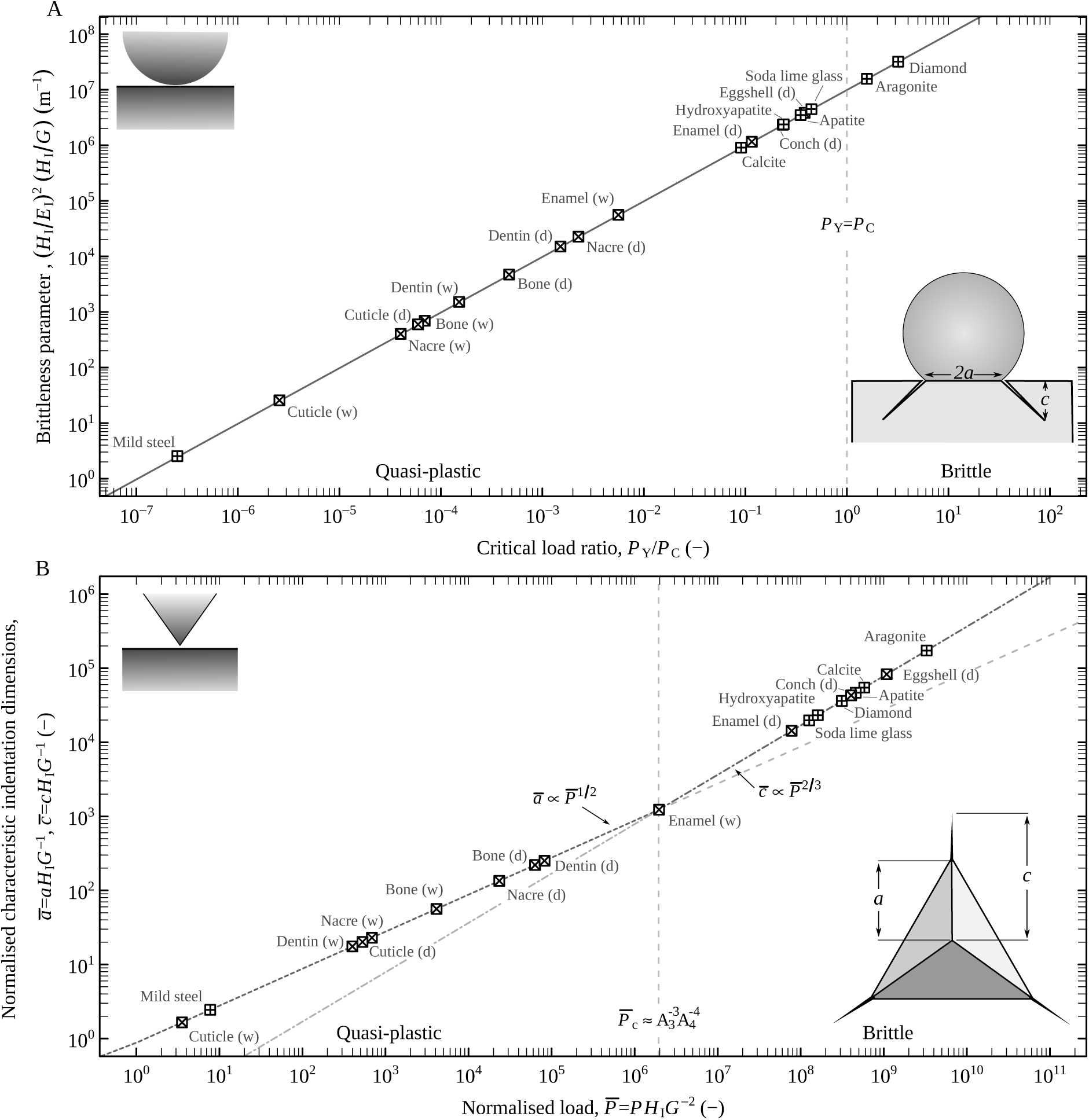
(A) Blunt contacts can cause damage *via* brittle fracture or quasi-plastic deformation. The dominating damage mode is set by a ‘brittleness parameter’, and the radius of curvature of the indenting body (here set to an arbitrary value of 1 mm). If the ratio between the load required to initiate quasi-plastic deformation and the load to induce a crack, *P*_Y_*/P*_C_, is larger than one, brittle fracture will dominate, and vice versa. For *R* < 4 mm, all biological materials included in this data set would be in the quasi-plastic regime. (B) If the indenter is sharp, yield is imminent. Cracks will only exceed the width of the residual impression resulting from quasi-plastic deformation once a critical load is exceeded (the normalised indentation dimensions shown here have been calculated using an arbitrary value of *P* = 1 N, and 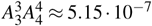 for a Berkovich indenter). For *P* < 5 mN, none of the biological materials included in this data set would crack. Increasing or decreasing the radius of the indenting particle (for (A)) or the indentation force (for (B)) will shift the materials to the right or left, respectively. The letters in brackets indicate if measurements were performed in hydrated *(w)* or dehydrated *(d)* conditions. Both plots illustrate the enourmous effect of both the composite design and hydration on biological materials (compare the data for pure aragonite to nacre, hydroxyapatite with dentin and enamel, and apatite with bone); both increase the resistance to fracture by orders of magnitude. The only exception appears to be chicken eggshell. Hydrated materials are less likely to crack, because of their increased toughness, and decreased indentation hardness and modulus. Data are from (21, 122, 123, own data) (biological materials), (121) (mild steel, soda lime glass, and diamond), (62, 124) (aragonite), (125) (apatite, calcite) and (126, 127) (hydroxyapatite).

In ideal brittle materials, the damage zone arising from fracture is characterised by the length of a coneor penny-shaped crack, *c*, which scales as

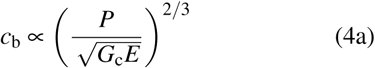

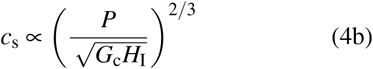

for blunt and sharp contacts, respectively (117, 118). Here, *G*_c_ = *T*^2^*/E* is the work of fracture, which is assumed to be independent of the crack length, and *E* is the Young’s modulus. The difference in the denominator, *E* vs. *H*_I_, arises because cracks in blunt contacts extend during loading, while radial cracks in sharp contacts grow to their final length during unloading, driven by a residual stress field 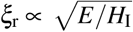 (119).

The damage zone induced by quasi-plasticity, in turn, is characterised by the depth of the residual impression, *h*_r_. The Oliver-Pharr model can be used to find

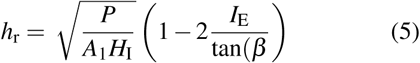

where *A*_1_ = *π*cotan^2^(*β*) is an indenter-dependent constant (*A*_1_ = 24.5 for a Berkovich tip). The term in the bracket represents the ratio between the measured contact pressure and the purely elastic limit. For a spherical indenter, the face inclination angle varies with contact depth as tan(*β*) ≈ *a/R* (120), where 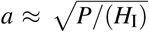 is the contact radius, so that

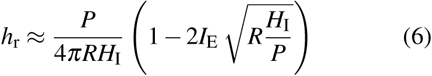

The second term reflects the fact that ‘yielding’ in blunt contacts will only occur at some finite load, *P*_Y_ > 4*H*_I_(*I*_E_)^2^*R*^2^, which is of identical form as the prediction derived by Rhee et al. (121), but with a somewhat larger constant (4 compared to 0.85, obtained by experimental calibration).

Unsurprisingly, a large work of fracture reduces the size of ensuing cracks, independent of whether the contacts are sharp or blunt. In addition, cracks are controlled by *E*_I_ in blunt contacts, and *H*_I_ in sharp contacts. Notably, the resistance to quasi-plastic deformation is not generally controlled by *H*_I_, but by the elasticity index, *I*_E_. Assuming that wear is proportional to the size of the damage zone, these simple guidelines can be used to rank biological materials by their resistance to mechanical wear (see table 1).

### The competition between quasi-plasticity and brittle fracture

While the previous arguments provide simple guidelines for ranking materials according to their ability to minimise the size of the damage zone(s) resulting from indentation, they do not address the question which of the damage modes is dominating, or indeed more likely to occur in the first place. In blunt contacts, materials can crack before they yield – a simple predictive index can be found by comparing the load required to initiate brittle fracture, *P*_C_, with the load required to cause quasi-plastic deformation, *P*_Y_ (121):

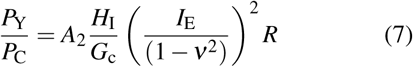

where *A*_2_ ≈ 10^−4^ is a dimensionless constant obtained by experimental calibration (note that Tabor’s law has been used in the derivation of eq. 7, see 121). If *P*_Y_*/P*_C_ > 1, brittle fracture will dominate, while quasi-plasticity will be the dominant damage mode if *P*_Y_*/P*_C_ < 1 (121). Equation 7 also includes the radius of the indenter, *R*, illustrating that quasi-plasticity, a volume effect, and brittle fracture, a surface effect, are linked by a characteristic length scale. This length scale, and therefore the propensity of a material to crack, is set by the ‘brittleness parameter’, 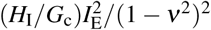 (121); large values indicate that small (but blunt) particles will likely induce a crack, whereas small values imply that the dominating damage mode will be quasi-plastic. The brittleness parameter is plotted against the load ratio for a number of biological and engineering materials in figure 4, assuming an arbitrary particle radius of 1 mm, and smearing the term containing Poisson’s ratio into the insecurity associated with the experimental constant *A*_2_. Even for such relatively large particles, biological materials are almost exclusively in the quasi-plastic regime, indicating that the ratio between indentation hardness and modulus is the parameters governing wear resistance.

If the wear-inducing body is sharp, yield occurs practically instantaneously, due to the divergent contact stresses. The competition between the two damage modes can then be assessed by comparing the width of the residual impression induced by quasi-plasticity, *a*_r_ (see inset in figure 4 b):

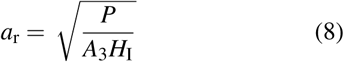

with the length of the crack introduced by brittle fracture, *c*, given by eq. 4b (including another geometrical constant, *A*_4_, in the nominator). Equating eqs. 8 and 4b yields an approximate prediction for the load at which *c* = *a*_r_, or, in other words, the load required to initiate a crack (see inset in figure 4 and 19, 128):

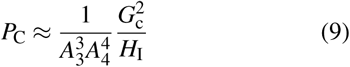

where *A*_3_ and *A*_4_ are constants relating to the indenter geometry. Equations 4b, 8 and 9 can be used to generate a universal plot of the normalised damage lengths against the normalised indentation load, which is shown in figure 4 B for an arbitrary load of 1 N, and a Berkovich indenter, for which 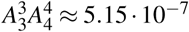 (for details on the normalisation, see refs. 19, 128).

While the available comparative data shown in figure 4 are limited, two conclusions emerge. First, quasi-plasticity is highly competitive in biological materials (see also ref. 21, for a similar argument): Cracks would be avoided altogether if *P* < 2mN (sharp contacts), or *R* < 3mm (blunt contacts). In contrast, shifting all investigated biological materials into the brittle fracture regime would require *P* > 0.6MN (sharp contacts), or *R* > 400m (blunt contacts). As a consequence, even relatively large wear particles will polish rather than grind biological materials. Second, the competitiveness of quasi-plasticity appears to be largely based on a composite design and on hydration (cf. hydrated and dehydrated nacre to pure aragonite, dentin and enamel to pure hydroxyapatite, and bone to pure apatite in figure 4). If these design features play any role in damage control, it is to avoid cracking, suggesting that quasi-plastic deformation may be less deleterious, or easier to control. The only exception to this rule appears to be chicken eggshell, which has a brittleness comparable to its main constituent calcite (we note that there has been some controversy about the toughness of eggshell, and other authors have reported higher values 123).

If the suggested dominance of quasi-plastic deformation is to reflect adaptation, the selective pressures that would favour it compared to brittle fracture need to be identified. A first insight can be gained from Griffith’s criterion, which relates the material’s strength, σ, to a characteristic flaw size, *c*_f_:

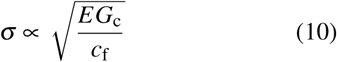

For sharp particles and loads smaller than *P*_C_, the relevant flaw size is *a*_r_ ∝ *P*^1/2^ (see eq. 8). For loads larger than *P*_C_, however, the relevant flaw size is *c* ∝ *P*^2/3^ (see eq. 4b), so that the degradation in strength due to flaws grows at a faster rate once brittle fracture becomes the dominating fracture mode *P*_C_ (129).

### The elasticity index as a measure of yield strength

One of the most well-known results of Tabor’s seminal work on the hardness of metals is that indentation hardness may be used as an index for yield strength, σ_Y_, because the ratio between the two is approximately constant, *H*_I_/σ_Y_ ≈ 3. Although used frequently, it remains unclear if this approximations is also valid for biological materials.

Tabor’s law can be rationalised *via* slip-line theory for ideally rigid-perfectly-plastic materials. However, Tabor’s law overestimates the constraint factor, ϒ = *H*_I_/σ_Y_ for materials that show considerable amounts of elastic deformation. Such plastic ideally-elastic solids can be analysed with expanding cavity models, which liken indentation to the expansion of a void driven by internal pressure. Yu and Blanchard (130) have proposed an expanding cavity model for elastic perfectly-plastic solids, based on the simplifying assumption that the material underneath the indenter will deform elastically until a critical pressure is reached, from where on the contact pressure is given by slip-line theory. In this model, the constraint factor ϒ = *H*_I_/σ_Y_ can be written as a function of the elasticity index, and the indenter geometry:

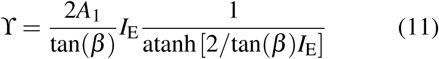

where *A*_1_ is a constant depending on the indenter geometry and the yield criterion (for a Berkovich tip and the von Mises yield criterion, *A*_1_ ≈ 2.75. See 130). For *I*_E_ = 0.05 and a Berkovich indenter, ϒ = 2.67, in reasonable agreement with Tabor’s relation.

The value of these predictions is limited by the lack of experimental data to put them to a test. Trim et al. (53) reported an elasticity index of *I*_E_ ≈ 0.055 for the dehydrated horn of bighorn sheep (*Ovis canadensis*), so that ϒ ≈ 2.66. The resulting prediction of *σ*_Y_ ≈ 72MPa is in reasonable agreement with the average yield strength measured in compression (*σ*_Y_ ≈ 65MPa, see ref. 53). The allegedly ‘strongest natural material’, the teeth of limpets, have a yield strength of approximately 2.25 GPa (estimated from figure 5 in 131). Using *E* = 120 GPa (131), *ν* = 0.22 (132), and eq. 11 yields *H*_I_ 6 GPa, which is in reasonable agreement with the available data for pure goethitie, but considerably larger than for complex goethitie (6.7 GPa and 1.1 GPa, respectively, see 132). Both examples suggest that the above simple model may be an acceptable approximation in some cases, and exploring this avenue further may be a valuable effort.

## Conclusions and outlook

Instrumented indentation has become the dominant tool for material characterisation on small scales. In biological materials, the primary use of indentation is to relate the measured properties to ultrastructure and function, which requires a detailed understanding of their mechanical significance. Indentation hardness is a hybrid property quantifying the resistance to both reversible and irreversible deformation, and is hence inherently linked to the elastic modulus. This link has been explored with a simple series elastic-plastic deformation model, which provided an expression for the ‘true’ hardness, a property independent of the modulus, facilitating a quantitative comparison of the resistance to irreversible deformation across and within structural biological materials. Indentation hardness and modulus appear to be linearly related in biological materials, and their ratio is an index for (i) the amount of relative irreversible deformation in the contact zone, (ii) the yield strength of the material, and (iii) specific types of mechanical damage. As quasi-plasticity is highly competitive with brittle fracture in biological materials, the ratio between indentation hardness and modulus can be used to rank materials by their damage resistance.

Two elephants in the room have thus far been largely ignored. First, we have repeatedly referred to ‘plasticity’ and ‘yield’, without providing any detail on what these phenomena actually involve. Both concepts are well understood in ‘classic’ engineering materials, but remain severely understudied in most biological materials (but see 133). Does the onset of yield involve dislocation motions and twinning along well-defined crystallographic planes, densification by compaction, phase transformations, micro-cracking, all of these or something completely different? Yield processes are unlikely to be material-invariant, so that the applicability of the previous arguments must be assessed from case to case. If yield induces microcracks, these can coalesce, in which case quasi-plasticity can be as deleterious for strength degradation, fatigue and wear resistance as brittle fracture (121), rendering it a particular problem for most biological materials. Second, most if not all biological materials are viscoelastic, so that the assumption of ideal elastic–plastic behaviour implicit in the previous analysis is unrealistic. However, comparative data on the viscoelastic properties of biological materials is almost completely absent. The data used in this study were dominantly collected with a trapezoidal load protocol, so that the measured modulus should be interpreted as a infinite-time indentation modulus (provided that the hold-period allowed sufficient time for relaxation or creep to converge). As a consequence, the estimated elasticity indices represent an upper bound. Viscoelasticity may be particularly important in damage control – as the residual stress field scales with *I*_E_*−*1, a time-dependent modulus can relieve some of these stresses into creep or relaxation, instead of feeding them into the extension or initiation of a crack. The immediate implication of this simple argument is that wear processes in biological materials must be rate-dependent in a complex fashion, but little research has addressed this phenomenon in detail.

Another hallmark of biological materials is that their mechanical properties are hydration-dependent. A meta-analysis of the available data underlined that hydration is particularly important in soft materials, and generally strongly reduces the propensity of materials to fracture, at the cost of making them more vulnerable to quasi-plastic deformation. The mechanistic basis for the influence of hydration on toughness, modulus, and hardness remains unclear, highlighting another important area of future research.

## Materials & Methods

Indentation modulus, hardness and fracture toughness of biological materials were collated from the literature. Relevant papers were identified with google scholar, using the search terms ‘biology+indentation’, ‘biology+hardness’, and ‘biological materials’. Data were either taken directly from the text or tables, or extracted from figures using the free tool *WebPlotDigitizer*, developed and maintained by Ankit Rohatgi (http://arohatgi.info/WebPlotDigitizer). All data can be found in the supplementary material.

## Acknowledgements

The authors would like to thank the Oyen-Lab and Jan-Henning Dirks for fruitful discussions and helpful comments on early versions of this manuscript. This study was supported by a Senior Research Fellowship from Clare College, Cambridge (to DL).

## Appendix

### Derivations

We make the simplifying assumption that any deformation during indentation is either irreversible on the time scale of the experiment (‘plastic’) or reversible (‘elastic’). The irreversible deformation is represented by a plastic component characterised by a ‘true’ hardness *H*, while the elastic deformation is represented by an elastic component characterised by a plane strain modulus *E′* = 1/(1 − *ν*^2^). The respective constitutive relationships between force and displacement are (14)

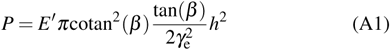

for the elastic component, and

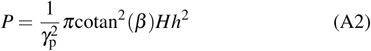

for the plastic component (*γ*_e_ and *γ*_p_ are constants relating to the depression or elevation of the contact perimeter. For elastic contacts, *γ*_e_ = *π/*2. See 14). These two components superimpose to determine the relationship between load and displacement during loading, which is of the form

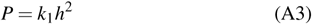

The load on two elements connected in series is identical to the total applied load, while the displacement is additive, so that from eqs. A1-A3 we find

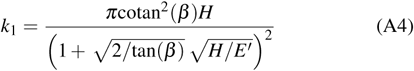

where we assumed *γ*_e_ = *γ*_p_ to ensure compatibility at maximum load. Using 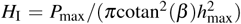 and eq. A3 yields eq. 1 in the main manuscript.

### Additional statistical analyses

#### The influence of indenter geometry on the relationship between indenation hardness and modulus

In order to remove the variation introduced by the different indenter geometries, eq. 2 was used to calculate the ‘true’ hardness, *H* for all materials, which was then rescaled into a Berkovich indentation hardness with eq. 1. A reduced major axis regression on log-transformed values yielded 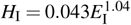 (95% confidence intervals (0.040; 0.047) and (1.02; 1.06), R^2^ 0.96), almost identical to the result obtained from the original data.

#### The sensitivity to hydration

The sensitivity to hydration decreases with hydrated indentation modulus as 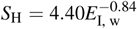 (R^2^ =64%, 95% CI (3.26; 5.92) and (-0.96; 0.74), respectively). A number of other models are summarised in table A1. Dehydration appears to have a stronger effect on indentation modulus compared to hardness for materials with small values of *H*_I, w_ and *E*_I, w_, while indentation hardness is more strongly affected in the limit of large *H*_I, w_ and *E*_I, w_ (based on the fitted parameters of a reduced major axis linear regression on log-transformed data, see table A1l). However, this statistical effect may be an artefact, first, because ‘wet’ contacts are likely more viscoelastic, impacting both indentation hardness and stiffness, but not necessarily in equal measures, second, because the conditions of the hydrated measurements differed between studies (some experiments were conducted in submersion, while others were conducted with ‘fresh’ specimen), and third, because re-hydration of previously dehydrated materials may not fully restore the initial material properties (60).

**Table A1:**
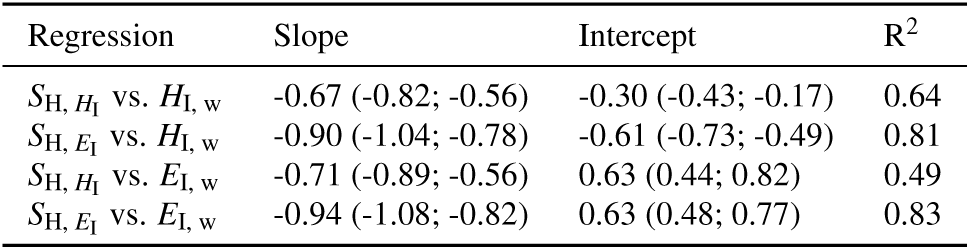
Results of reduced major axis regressions on logtransformed data, with wet indenation hardness and modulus in GPa. The values in brackets indicate 95% confidence intervalls.

## Footnotes

1 The distinction between sharp and blunt contacts is somewhat arbitrary – sharp indenters are those with a steep angle of the tangent at the point of contact. Irreversible deformation will necessarily precede cracking if the contact is sharp, while the deformation can remain dominantly elastic prior to fracture if the indenter is blunt. To a first approximation, spheres are blunt, and conical, Vickers, Berkovich and cube corner indenters are sharp.

